# A ride in the park: Cycling in different outdoor environments modulates the auditory evoked potentials

**DOI:** 10.1101/455394

**Authors:** Joanna E. M. Scanlon, Eden X. Redman, Jonathan W. P. Kuziek, Kyle E. Mathewson

## Abstract

In this study, we investigated the effect of environmental sounds on ERPs during an auditory task, by having participants perform the same dual task in two different outdoor environments. Participants performed an auditory oddball task while cycling outside both in a quiet park and near a noisy roadway. While biking near the roadway, an increased N1 amplitude was observed when evoked by both standard and target tones. This may be due to attentional processes of enhancing sound processing in the noisier environment. No behavioural differences were found. Future directions include investigating auditory ERPs in more realistic studies outside of laboratory.

## 1. Introduction

Cognitive neuroscience traditionally requires subjects to sit in highly controlled environments, and avoid any kind of natural light, sound, and movement during recording. This is because all uncontrolled sensations have the possibility of introducing noise into the EEG signal (Schlögl, Anderer, Roberts, Pregenzer, & Pfurtscheller, 1999; White & Van Cott, 2010), and this noise can be a large factor in the statistical power when analyzing EEG and ERP data (Luck, 2014). New technologies are beginning to make it easier to record laboratory-quality EEG data outside of the lab while subjects are mobile (Debener et al., 2012; Kuziek, Sheinh, & Mathewson, 2017; Scanlon et al. 2017b). Performing experiments in real-world environments allows greater generalizability and understanding of how the brain works in everyday life. While the effects of movement on EEG data are beginning to be understood, few studies have attempted to investigate the effect that different environments may have on the way the brain functions. The present study was conducted in a fully mobile, outdoor environment, in order to investigate the ways in which different outdoor environments may affect the ERP.

In mobile EEG literature, the oddball task is commonly used due to its high signal to noise ratio in the detection of the P3, as well as for the ability to infer changes in attentional resources due to a concurrent task (Polich, 1987; Polich & Kok., 1995). This makes the oddball task particularly effective for learning about the way one’s environment can affect attention, especially when used as a dual-task paradigm with movements such as walking. Debener et al. (2012) recorded brain activity using a small consumer passive-electrode wireless EEG mounted to a typical laboratory electrode cap, and had participants perform an auditory oddball task while either walking outdoors or sitting indoors. The authors demonstrated that the P3 was significantly larger when generated while sitting indoors than during outdoor walking. Using a similar wireless EEG system, de Vos, Kroesen, Emkes, & Debener (2014) found a significantly larger P3 response to a three-stimulus oddball paradigm while participants were sitting compared to walking outside. Zink, Hunyadi, Van Huffel, and de Vos (2016) used a passive wireless mobile EEG system and asked participants to perform an auditory oddball paradigm during mobile cycling, stationary pedaling and sitting in a natural outdoor environment. The authors found no difference in root mean square (RMS) data noise or P3 amplitude between sitting and pedaling, but demonstrated a significant increase in noise and near-significant decrease in P3 amplitude while participants were cycling. Another study by Ladouce, Donaldson, Dudchenko, & Ietswaart (2019) demonstrated a reduced P3 while walking as well as being simply wheeled around in comparison to sitting, indicating that the effect of walking on the P3 is at least partly due to visual influences on attention. Additionally, this can be seen in other navigation scenarios. Chan, Nyazika & Singhal showed reduced P3 amplitude while participants drove in a driving simulator with a passenger, compared to rest. Scanlon, Townsend, Cormier, Kuziek, & Mathewson (2017b) had participants perform an oddball task both while cycling outside and sitting inside, using a regular non-mobile EEG system with active electrodes and optimized for mobility using a tablet computer. Consistent with other mobile EEG studies mentioned we found a significantly decreased P3 amplitude while cycling outside (Scanlon et al., 2017b). Altogether these studies indicate that the P3 to an oddball task is reduced during concurrent purposeful movement.

Our research group at the University of Alberta has found cycling to be an ideal task for mobile EEG research, as it provides a rich motor task with smooth, straightforward movement while navigating the environment. Our first cycling study looked at the effects of cycling on EEG data, and had participants perform an auditory oddball task before, during and after a sub-aerobic stationary cycling task (Scanlon et al., 2017a). We found increased EEG data noise during the stationary cycling task, however despite this noise, ERP signals were reliably recorded, with no ERP component differences between the cycling conditions. In a follow-up to this study (Scanlon et al., 2017b) we used compact technologies that could be fit inside a backpack, in order to have participants perform the auditory oddball task while cycling outside. Participants were asked to listen to the oddball task through headphones both while cycling outside and while sitting in the laboratory. Baseline data noise was increased during the cycling condition; however, we were able to collect laboratory quality ERPs in both conditions. We demonstrated significantly decreased P3 amplitude and alpha band oscillations while participants cycled outside. This study also found an unexpected increase in the N1 component and decrease in the P2 component of the ERP, during both standard and target tones while participants were cycling outside. In a follow-up study (Scanlon, Cormier, Townsend, Kuziek, Mathewson, 2019) we investigated this effect further by having participants perform the same headphone auditory oddball task inside the lab with different sounds playing in the background of the task. Here, we found that we were able to replicate the N1 and P2 effects while playing outdoor noises in the background of the auditory oddball task, indicating that the effect may have been due to a selective attention process of filtering background noises in order to perform the task. These studies indicate that bringing EEG into ecologically valid environments opens up the possibility of demonstrating the way mechanisms such as auditory selective attention can be observed in real-world environments.

The current study builds on the findings of Scanlon et al., (2017b), and Scanlon, et al., (2019), using methods and analysis established in these studies. It is of particular interest to examine variations in ERP components between path cycling in quiet and noisy outdoor areas. If the N1 and P2 are in fact related to a selective attention process of filtering background noises in order to pay attention to task-relevant sounds, we would expect to see an increased N1 and decreased P2 while participants perform the same task in a noisier environment. For this experiment, each participant completed four 6-minute blocks in an auditory oddball task while both sub-aerobically cycling near a noisy roadway and in a quiet city park. Similar to Scanlon, et al. (2017b), active low-impedance electrodes were used. ERP magnitude, morphology and topography for the P3, N1 and P2 were analyzed, as well as levels of data noise and power spectra. Our first hypothesis is that due to increased ambient noise in the roadway condition, there will be an increased N1 and decreased P2 while participants cycle near the roadway. Our second hypothesis is that, similar to previous studies using ambient noise to manipulate N1 and P2 amplitude, there will be no behavioural effects between conditions.

## 2. Materials and Methods

### 2.1 Participants

15 people who were part of the university community participated in the experiment, however 5 were excluded due to technical issues during the recording, leaving 10 participants for final analysis (mean age = 23.4; Age range = 20-31; 4 female). Each participant received an honorarium of $20. All participants had normal or corrected to normal vision and with no history of neurological or hearing problems. Experimental procedures used were approved by the Internal Research Ethics Board of the University of Alberta.

### 2.2 Materials

Prior to the experiment, participants selected one of two bicycles (Kona Mahuna), which differed only in frame size (17 or 19 inches) based on the participant’s height. Seat height was adjusted according to the participants’ comfort level. Bicycle gears were set to the 2^nd^ gear in both the front and back derailleurs, in order to allow participants to pedal constantly and evenly throughout the trials at a sub-aerobic level. Data was collected using a Microsoft Surface Pro 3 running Brain Vision recorder (Brain Products), and powered by an Anker Astro Pro2 20000mAh Multi-Voltage External Battery or or Tzumi Extreme PocketJuice 6000mAh battery. Due to technical problems with the Anker astro battery, the Tzumi battery was used for 8 subjects. The various technologies mentioned were connected using a Vantec 4-Port USB 3.0 Hub.

To run the oddball task and mark the EEG data for ERP averaging, we used a Raspberry Pi 2 model B computer, running version 3.18 of the Raspian Wheezy operating system, using version 0.24.7 of the OpenSesame software (Mathȏt, Schreij, & Theeuwes, 2012; see Kuziek, Sheinh, & Mathewson, 2017 for validation study). A 900 MHz quad-core ARM Cortex-A7 CPU was connected through a 3.5 audio connector for audio output. To mark the data for EEG averaging, 8-bit TTL pulses were sent to the amplifier through a parallel port connected to the General Purpose Input/Output (GPIO) pins. A button connected to the Raspberry Pi was affixed to the handlebar of the bike, which was used both as a start button for each block, and a response button to the target tones. Participants wore a two-pocket backpack (Lululemon; Figure 1A, C) containing all of the stimulus generating and recording materials. In the *roadway* condition, participants rode along approximately 0.65 km of a shared use path next to River Valley Road in Edmonton, Canada (Figure 1B green). During the *park* condition, participants rode along approximately 0.65 km of a shared use path in MacKinnon Ravine Park, also in Edmonton, Canada (Figure 1B, blue). Both conditions were performed on the same day, within a 1-2 hour period, and condition order was randomly varied across participants.

**Figure 1.**
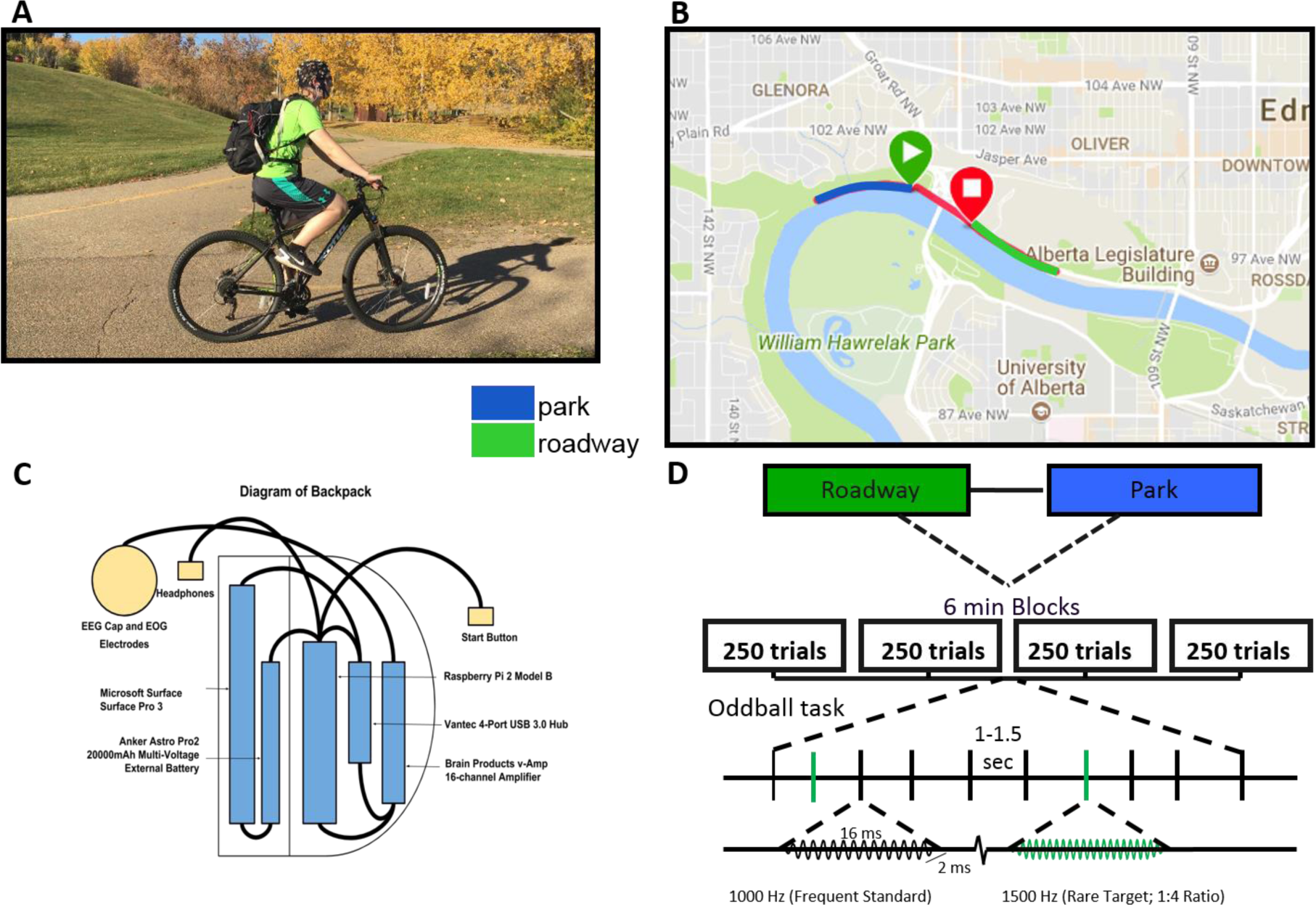
Biking apparatus and procedure for mobile EEG. A: The task was performed while cycling on a bicycle, with the participant wearing an EEG cap and backpack which contained the EEG equipment. B: A map of the cycling routes. C: Schematic layout of the contents of the mobile EEG backpack. The backpack had two pockets and contained the following: Anker Astro Pro2 20,000 mAh Multi-Voltage external battery or Tzumi Extreme PocketJuice 6000mAh battery, Raspberry Pi 2 model B, Microsoft Surface Pro 3, Vantec 4-Port USB 3.0 hub, Brain Products v-Amp 16 channel amplifier. D: During both cycling by the roadway and in the park, participants performed four 6 minute blocks of an auditory oddball task, with self-paced breaks in between. Within the oddball task, 20% of the tones were rare high-pitched tones (1500 Hz) and 80% were frequent low-pitched tones (1000 Hz). Each tone played for 16 ms with a 2 ms ramp-up and down. Tones were spaced 1-1.5 seconds apart.

Originally, a Garmin (Vivoactive HR) watch was used to monitor heart rate and speed during both conditions, however upon viewing this data, we found that the heart rate data could not be accurately used due to inconsistent connection to the skin in most participants. However, speed data was used to estimate an average moving speed over the whole experiment of 6.9 km/hr.

### 2.3 EEG Recording

Active wet electrodes were selected for EEG recording in this study based on previous studies by Laszlo et al. (2014) comparing active and passive amplification electrodes at differing levels of impedance, as well as by Oliviera et al. (2016) who compared active wet, passive wet and passive dry electrodes during a mobile task. Both of these studies demonstrated that active wet electrodes afford cleaner and better quality signals (operationalized by lower data noise and higher signal-to-noise ratio) in non-ideal recording conditions. Fifteen Ag/AgCl pin electrodes were arranged in 10-20 positions (Fp2, F3, Fz, F4, T7, C3, Cz, C4, T8, P7, P3, Pz, P4, P8, and Oz). Additionally, a ground electrode was embedded in the cap at the Fpz position, along with two reference electrodes clipped to the right and left ears. To lower electrode impedance and decrease the effect of sweat on the data, SuperVisc electrolyte gel was used, followed by mild abrading of the skin with a blunted syringe tip (Kappenman & Luck, 2010; Luck, 2005; Picton & Hillyard, 1972). The aforementioned techniques and gel application continued until all the electrodes had impedance levels lower than 10 kΩ, as measured using an impedance measurement box (BrainProducts) and until the raw data appeared to be clean and free of excessive noise. Between conditions, a second impedance check was performed and any problematic electrodes were corrected. In addition to the 15 EEG electrodes, two reference electrodes and one ground electrode, horizontal and vertical bipolar electrooculogram (EOG) was recorded through passive Ag/AgCl easycap disk electrodes placed above and below the left eye, as well as 1 cm from the outer canthus of each eye. These passive electrodes were used for the EOG because the AUX ports of our amplifier do not support active electrodes and we did not see a benefit to giving up 4 EEG channels for active-wet EOG sensors. These EOG electrodes can nonetheless record reliable signals due to their placement on the face, where there is generally less hair to impede the signal. Dirt and oils at the EOG electrode sites are removed using NuPrep preparation gel, followed by cleaning the skin with an antibacterial sanitizing wipe prior to electrode placement in order to reduce impedance levels of the EOG electrodes based on visual inspection of the raw data. These bipolar channels were recorded using a pair of BIP2AUX converters plugged into the AUX ports of the V-amp amplifier, along with a separate ground electrode placed on the central forehead. EEG and EOG was recorded using a V-amp 16 channel amplifier (Brain Products) through a laptop USB port. Data were digitized at 500 Hz with a 24 bit resolution. The data were filtered using an online bandpass with 0.1 and 30 Hz cutoffs, along with a 60 Hz notch filter. These narrow filters were recommended in the actiCap Xpress manual in order to increase signal quality and reliability in mobile settings (Brain Products, 2014).

Data collection was completed between the months of August and November. When outside, participants sometimes wore a toque (knitted cap) over the EEG cap to prevent the electrolyte gel from drying out, goggles to prevent excessive eye blinking during the trials, and gloves, if desires due to the weather (Temperature range = −4 - 17°C; Mean temperature = 11°C). During each 6 minute block the participant would follow a research assistant on a leading bike, and travel approximately 650 meters from the starting point until the tones stopped. After which they would turn around, have the data visually inspected for noise and electrode problems by the research assistant, and complete another 6 minute block going in the opposite direction.

Participants were instructed to stare straight ahead while following the lead experimenter, and pedal slowly and continuously, in order to maintain mobility at a sub-aerobic level.

### 2.4 EEG Analysis

Analyses of EEG data were completed using MATLAB 2016b with EEGLAB (Delorme & Makeig, 2004), with custom scripts. Following data recording, the EEG data was re-referenced to an average of the left and right ear lobe electrodes (Figure 2). TTL pulses time-locked to the onset of target and standard tones were marked in the EEG data during recording, and were then used to construct epochs 1200-ms long (including a 200-ms pretrial baseline).

**Figure 2.**
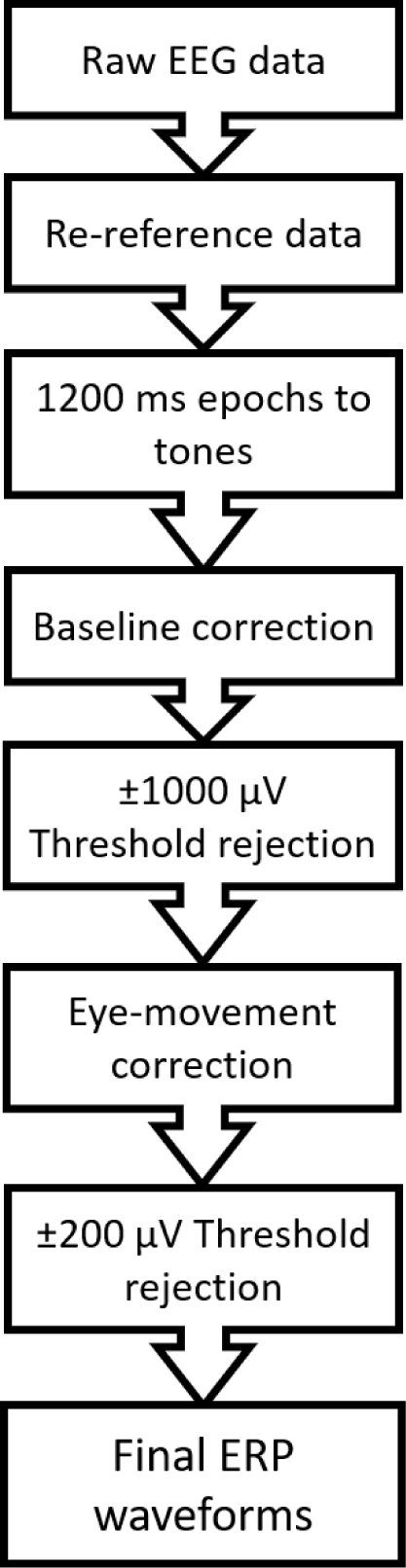
ERP analysis methods, in logical order from raw EEG data to final ERP waveforms.

Average voltage in the first 200-ms baseline was subtracted from the data for each trial and each electrode. To remove artifacts caused by movement, saturated signals, amplifier blocking, and any other non-physiological factors that may affect eye movement correction, any trials in either of the conditions with a voltage difference from baseline larger than ±1000 μV on any channel (including eye channels) were removed from further analysis. Then a regression-based procedure was used to correct for eye movements (details in the following paragraph). Following eye-movement correction, a second threshold was applied to remove any trials with over ±200uV voltage difference from baseline. After this procedure, over 88% of trials were retained in each condition. Overall, artifact rejection left approximately equal trial numbers per participant in the park (M_targ_ = 183; range_targ_ = 127-200; M_stand_ = 722; range_stand_ = 457-800; SD = Standard Deviation) and roadway (M_targ_ = 179; range_targ_ = 124-200; M_stand_ = 708; range_stand_ = 498-800) conditions, which we then used for the remaining analysis.

A regression-based procedure was used to estimate and remove variance due to EEG artifacts caused by blinks, as well as horizontal and vertical eye movements (Gratton, Coles, & Donchin, 1983). This technique uses a template-based approach to identify blinks, then computes propagation factors such as regression coefficients, and predicts the vertical and horizontal eye channel data from each electrode’s signal. Then, weighted by these propagation factors, this eye channel data is then subtracted from each channel, allowing us to remove most variance in the EEG data that can be predicted by movements from the eyes. In order to keep as many trials as possible for both conditions, no further filtering or rejection was done on the data.

### *2.6* Statistical analysis

In this study, we intend to analyze behavioural responses, ERPs and spectral power between the two conditions. Behavioural analysis consists of pair-wise t-tests between average oddball task response times and response accuracies in the two conditions. ERP analysis for this study will focus on the N1 (Fz and Pz; 100-175 ms), P2 (Fz and Pz; 175-275 ms), and P3 (Pz; 334-434 ms) ERP components. These electrodes and time windows were chosen to surround clear peaks in the topography and data present in both conditions, and based particularly on previous comparable literature (Scanlon et al., 2017a; Scanlon et al., 2017b; Scanlon et al., 2019). To assess both condition differences and differences between standards and targets, a 2 × 2 repeated measures ANOVA was conducted for the mean amplitude within the time window for each component with the first factor being standards vs. targets and the second being the two location conditions. Where significant group differences were found between conditions, pair-wise Bonferroni corrected t-tests were then used to further investigate how the conditions differed. Additionally, we calculated Cohen’s d to estimate effect sizes, as well as 95% confidence intervals for all t-tests.

As the task used a button press for response to target tones, it is possible that effects of muscular preparation and activation may be included in the ERP and oscillation effects for targets. Additionally, there is no reason to predict differences between standard and target tones in prestimulus data noise. Therefore, only standard tones were included in the spectral and RMS analysis. For single-trial and baseline ERP RMS data noise a Wilcoxon Signed rank test was used to measure differences between the conditions. Mean differences and 95% confidence intervals were obtained using the Hodges-Lehmann estimator. Pair-wise t-tests were used to test for spectral differences between conditions.

## 3. Results

### 3.1 Behavioural analysis

In order to assess behavioural effects in this task, we tested for condition differences in response accuracy and average reaction time to the target stimuli in the task, depicted in Figure 3. A two-tailed paired samples t-test indicated no significant difference in response accuracy between the two conditions (M_diff_=-1.60; SD_diff_=2.49; t(9) =-2.03; p=0.0729; *Cohen’s d* = −0.642; CI_95%_ = −3.38-0.183). The same test was used to test for differences in average reaction time, finding no differences between conditions (M_diff_=4.10; SD_diff_=26.2; t(9) =0.495; p=0.633; *Cohen’s d* = 0.157; CI_95%_ = −14.7-22.9).

**Figure 3.**
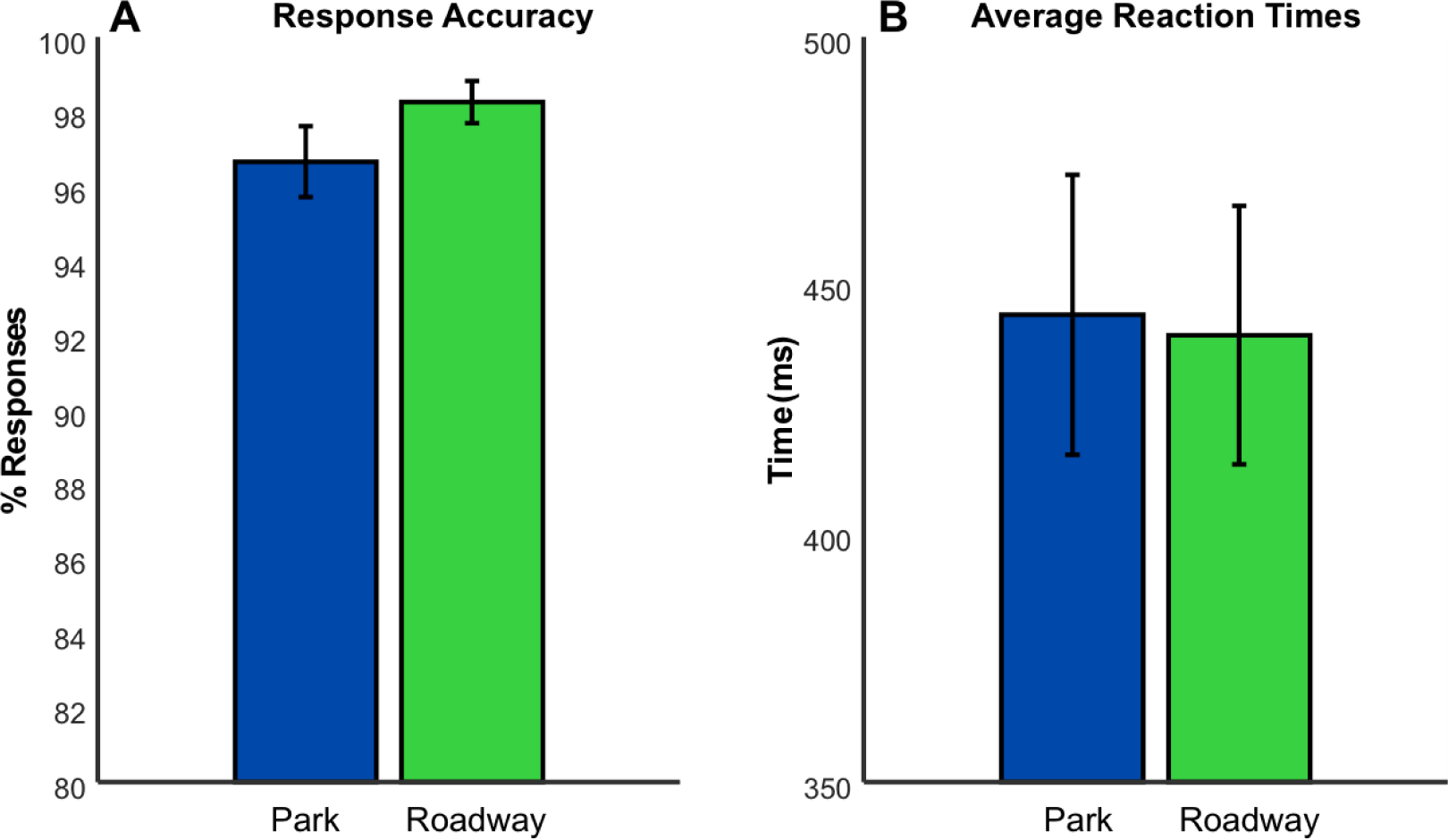
Behavioural analysis. A: Bar graph depicting mean and within-subjects standard error for the percentage of targets that received a response within 2 seconds after the tone onset across participants for both conditions. B: Bar graph depicting the mean and within-subjects standard error for the average time taken to respond (with missed responses removed) across participants for each condition.

### 3.2 Spectral analysis

We used two separate methods to estimate the data noise on individual trials. First, we took an average of the spectra over each EEG epoch in the Fz and Pz electrode locations. Each participant’s data was randomly sampled for 290 of their artefact-removed standard trials. A fast Fourier transform (FFT) was then calculated through a procedure of symmetrically padding the 600 time point epochs with zeros, making for each epoch a 1,024-point time series, which provided frequency bins of .488 Hz. Because the data were collected online with a low-pass filter of 30Hz, only frequencies measuring up to 30 Hz were plotted. We then calculated the spectra for each participant by computing the average of the 290 spectra for each participant, and then combining these into the grand average spectra, as shown in Figure 4A for the Fz and Pz channels. Shaded regions depict the within-subjects standard error of the mean across participants. Evident in the plots, both conditions depict the expected 1/frequency structure in the data (Cohen, 2014). Also evident is a significant increase in beta oscillations (15-30Hz) at the Fz electrode location (M_diff_=0.0641; SD_diff_=0.07322; t(9)= 2.77; p=0.0218; *Cohen’s d* = 0.851; CI_95%_ = 0.0135-0.156) during the park compared to the roadway condition, as tested with a two-tailed paired samples *t*-test.

**Figure 4.**
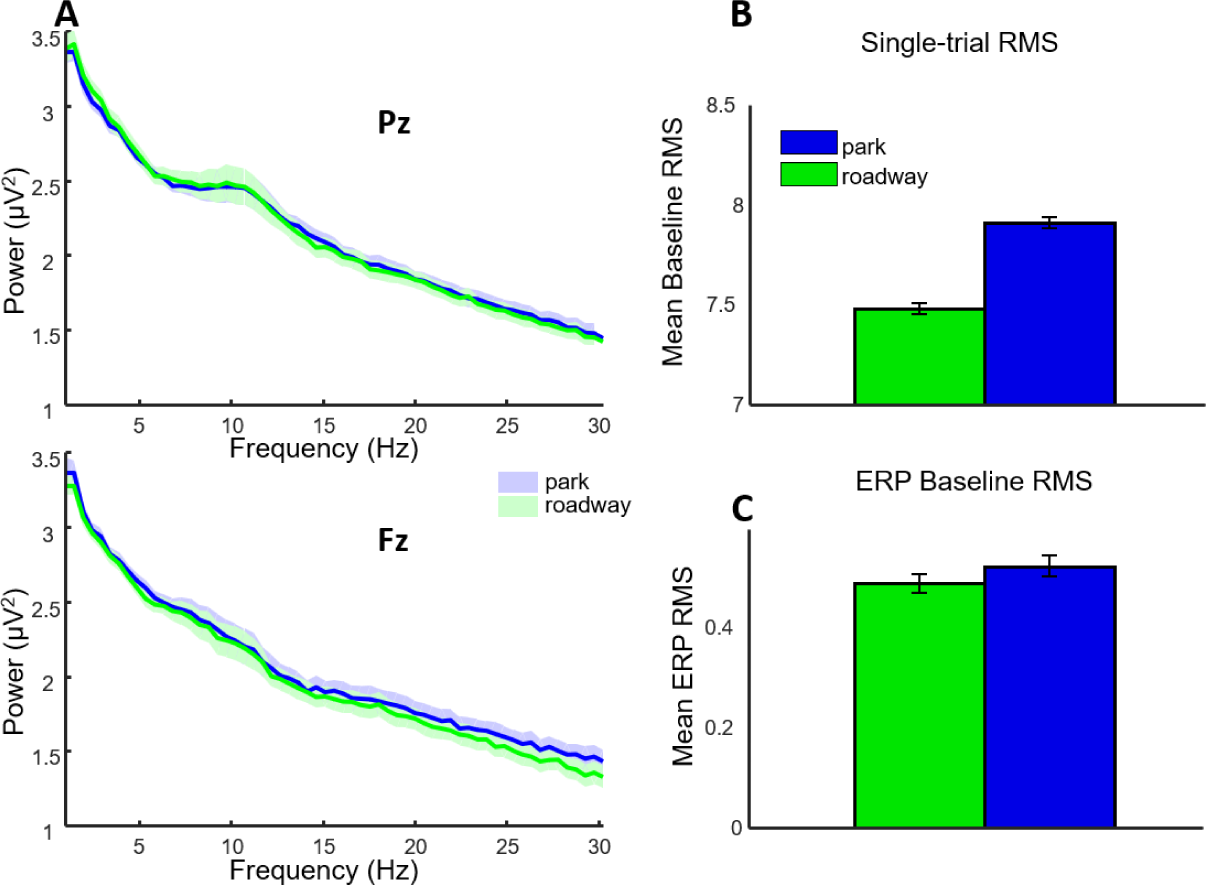
Spectra and data noise levels. A: Single-trial EEG spectra from electrodes Pz and Fz. B: Bar graph of average single-trial root mean square (RMS) grand average values. Error bars depict standard deviation of the permuted distributions. C: Bar graph of average ERP baseline RMS values. Error bars depict standard deviation of the permuted distributions.

### 3.3 Single-Trial and ERP Baseline Noise

As an additional estimate of the single-trial EEG noise, we computed the root mean square (RMS) value for the 200 ms baseline period for each standard trial (de Vos, Gandras & Debener, 2014). RMS is defined as the square root of the mean of squares for a series of numbers, and is equivalent to the average absolute voltage difference around the baseline, making this measure a good estimate of single trial noise in EEG data. Data were averaged over all electrodes for each trial, then over trials, then subjects. To avoid any interference from the evoked ERP activity, RMS values were taken from the baseline period, consisting of the 200 ms (100 time points) prior to each tone’s onset. To estimate an RMS distribution for each subject in the dataset, we used a permutation test that selects a different set of 290 epochs without replacement for each participant on each of 10,000 permutations before running second order statistics (Laszlo et al., 2014; Mathewson et al., 2017). A grand average single-trial RMS was calculated and recorded for each permutation and then condition, demonstrated in figure 4B. The park condition (M_RMS-EEG_ = 7.91; SD_RMS-EEG_ = 0.0288) had significantly larger single-trial noise levels compared to the roadway condition (M_RMS-EEG_ = 7.48; SD_RMS-EEG_ = 0.0272; M_diff_ = −0.432; CI_95%_ = −0.433 - −0.431; Wilcoxon signed rank test: z = 86.6; p < 0.0001).

We ran a similar permutation analysis within trial averaged ERPs, to test for noise that was not effectively averaged out across trials. Here, we used another permutation test of the 200 ms baseline RMS values to quantify the amount of noise in the participant average ERPs. This analysis is complementary to the above single-trial analysis, as the computation estimates the amount of phase-locked EEG data noise which is not averaged out over the trial with respect to tone onset. The computation randomly selected and averaged 290 of each participant’s standard artefact-removed trials without replacement. The obtained RMS values were then averaged over EEG electrode channels to create a grand average of all participants. This made 10,000 permutations after averaging over all participants, which was then used to compute second-order statistics (Figure 4C). The park condition (M_RMS-ERP_ = 0.522; SD_RMS-ERP_ = 0.0206) had a higher RMS value compared to the roadway conditions (M_RMS-ERP_ = 0.488; SD_RMS-ERP_ = 0.0193; M_diff_ = −0.345; CI_95%_ = −0.0351--0.0339; Wilcoxon signed rank test: z = 79.3; p < 0.0001).

### 3.4 ERP Morphology and Topography

Grand average ERPs calculated from each participant’s corrected and artefact-removed standard and target tones at electrode Pz are depicted in Figure 5A. Similar error levels can be observed within the two conditions. Evident from the plots is the expected P3 amplitude increase during target trials in the posterior topographical locations (Pz; Figure 5B). The expected oddball difference in the P3 is shown with an increased positive voltage between 334 and 434 ms following the infrequent target tones, compared with the common standard tones. We used this time window for all further analysis of the P3.

**Figure 5.**
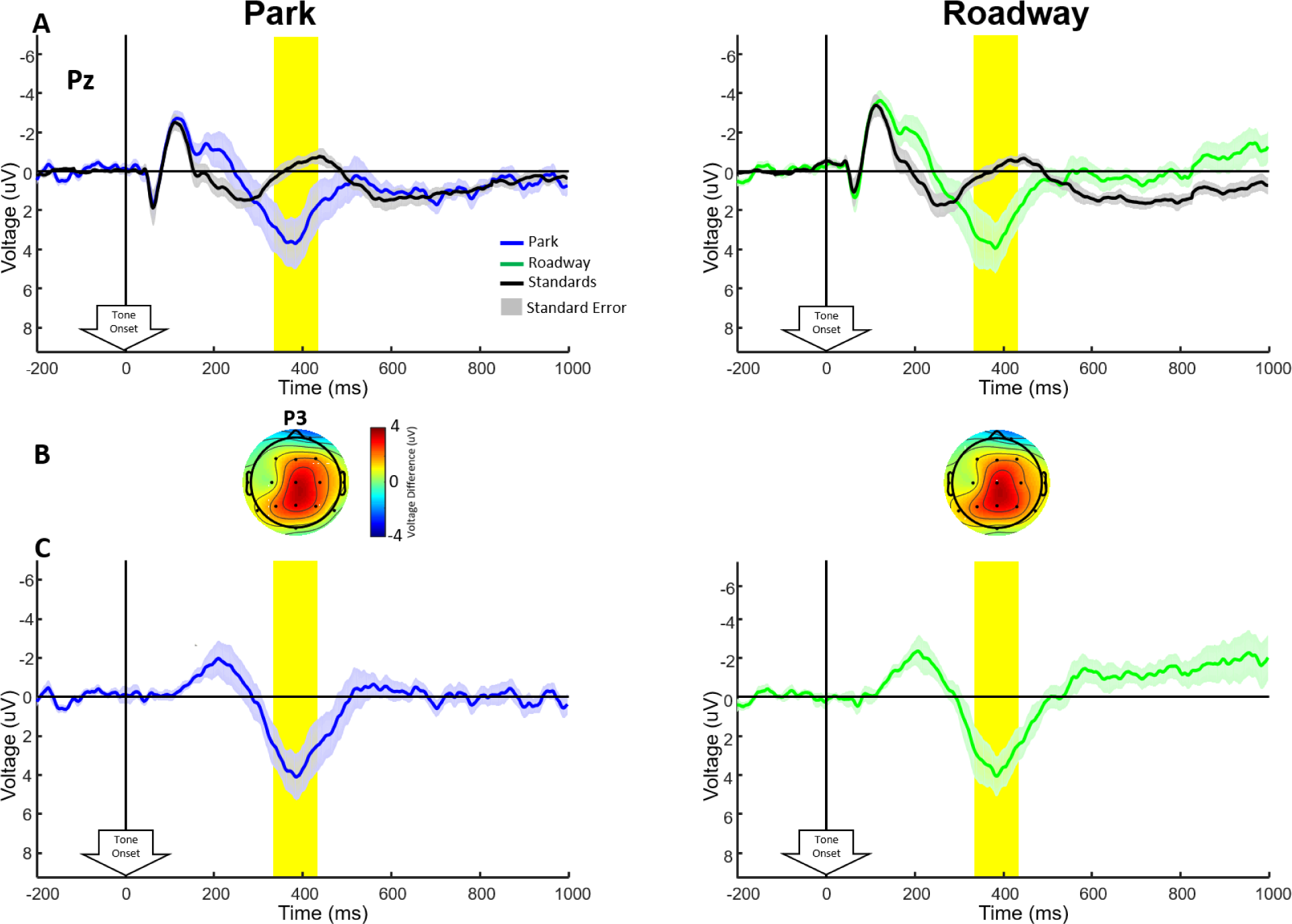
ERP grand averages. A: Grand-average ERPs computed at electrode Pz for all artefact-removed and eye movement corrected trials, for both standard (black) and target (colour) tones. B: Scalp topographies for grand-average ERP difference between standard and target tones in the P3 time window (indicated in yellow). C: ERP difference wave from electrode Pz for both conditions, with shaded regions depicting within-subjects standard error of the mean for this difference, with between-subjects differences removed (Loftus & Masson, 1994). Yellow highlighted regions represent the time window for the P3 analysis as well as topographic plots.

Figure 5B shows topographies of these target-standard differences within the P3 time window. Figure 5C plots the ERP difference waves at electrode Pz, which are created by subtracting the standard tone ERPs from target tone ERPs for each subject.

### 3.5 ERP Differences

Figure 5C shows difference waves plotted for both conditions at electrode Pz. Evident from the graphs is no significant P3 differences between the two conditions at Pz. To investigate stimulus and condition differences for the P3, a 2 × 2 repeated measures ANOVA test was carried out with the first factor being the stimulus (e.g. standards vs. targets) and the second factor being the condition (e.g. park vs. roadway). A main effect of stimulus was found for the P3(*F*(1,9) = 10.11, *p* = 0.0112, η_p_ = 0.529), with no main effect of condition (*F*(1,9) = 0.299, *p* = 0.598, η_p_ = 0.032) or interaction (*F*(1,9) = 0.0012, *p* = 0.973, η_p_ = 0.00). To further investigate the stimulus effects due to the oddball task, Bonferroni corrected (α = 0.025) two-tailed paired t-tests of these differences were performed within each condition. The target-standard differences were significant within both conditions (Park: M_diff_ = 3.35; SD_diff_ = 3.54; *t*(9) = 2.99; *p* = 0.015; *Cohen’s d* = 0.946; CI_95%_: 0.816-5.89; Traffic: M_diff_ = 3.37; SD_diff_ = 3.33; *t*(9) = 3.20; *p* = 0.011; *Cohen’s d* = 1.01; CI_95%_: 0.984-5.75)

### 3.6 N1 and P2 Amplitudes

In order to observe the effects of two different environments on general stimulus processing, grand averaged ERPs with a comparison of park vs. roadway at the Fz and Pz electrode locations are plotted in Figure 6A and B, respectively. A visual inspection of these plots shows increased amplitude in the N1 component in the roadway condition, both in targets and standards and at electrodes Fz and Pz. Another 2 × 2 repeated measures ANOVA test was carried out at each electrode location, showing a significant effect of stimulus (Fz: *F*(1,9) = 19.6, *p* = 0.00170, η_p_ = 0.685; Pz: *F*(1,9) = 26.0, *p* = 6.45e-04, η_p_ = 0.743) with larger (negative) amplitudes for the target tones (Table 1). A significant effect was also found for condition (Fz: *F*(1,9) = 6.41, *p* = 0.0321, η_p_ = 0.416; Pz: *F*(1,9) = 9.38, *p* = 0.0135, η_p_ = 0.510), but not for the interaction (Fz: *F*(1,9) = 0.478, *p* = 0.507, η_p_ = 0.05; Pz: *F*(1,9) = 0.406, *p* = 0.540, η_p_ = 0.0430). To further investigate the condition effects, two Bonferroni corrected (α = 0.025) two-tailed paired t-tests of these differences were performed within both standard and target stimuli at each electrode. The differences were not significant at Fz (Standards: M_dif f_ = 0.921; SD_dif f_ = 1.14; *t*(9) = 2.55; *p*=0.0310; *Cohen’s d* = 0.807; CI_95%_: 0.105-1.74; Targets: M_diff_ = 1.08; SD_diff_ = 1.43; *t*(9) = 2.37; p = 0.0417; *Cohen’s d* = 0.750; CI_95%_: 0.0501-2.10) but were significant at Pz (Standards: M_diff_ = 0.876; SD_diff_ = 1.00; *t*(9) = 2.77; *p* = 0.0217; *Cohen’s d* = 0.876; CI_95%_: 0.161-1.59; Targets: M_diff_ = 1.04; SD_diff_ = 1.14; *t*(9) = 2.88; *p* = 0.0180; *Cohen’s d* = 0.912; CI_95%_: 0.225-1.86). Figure 6C plots topographies of these differences, indicating a frontocentral distribution.

**Figure 6.**
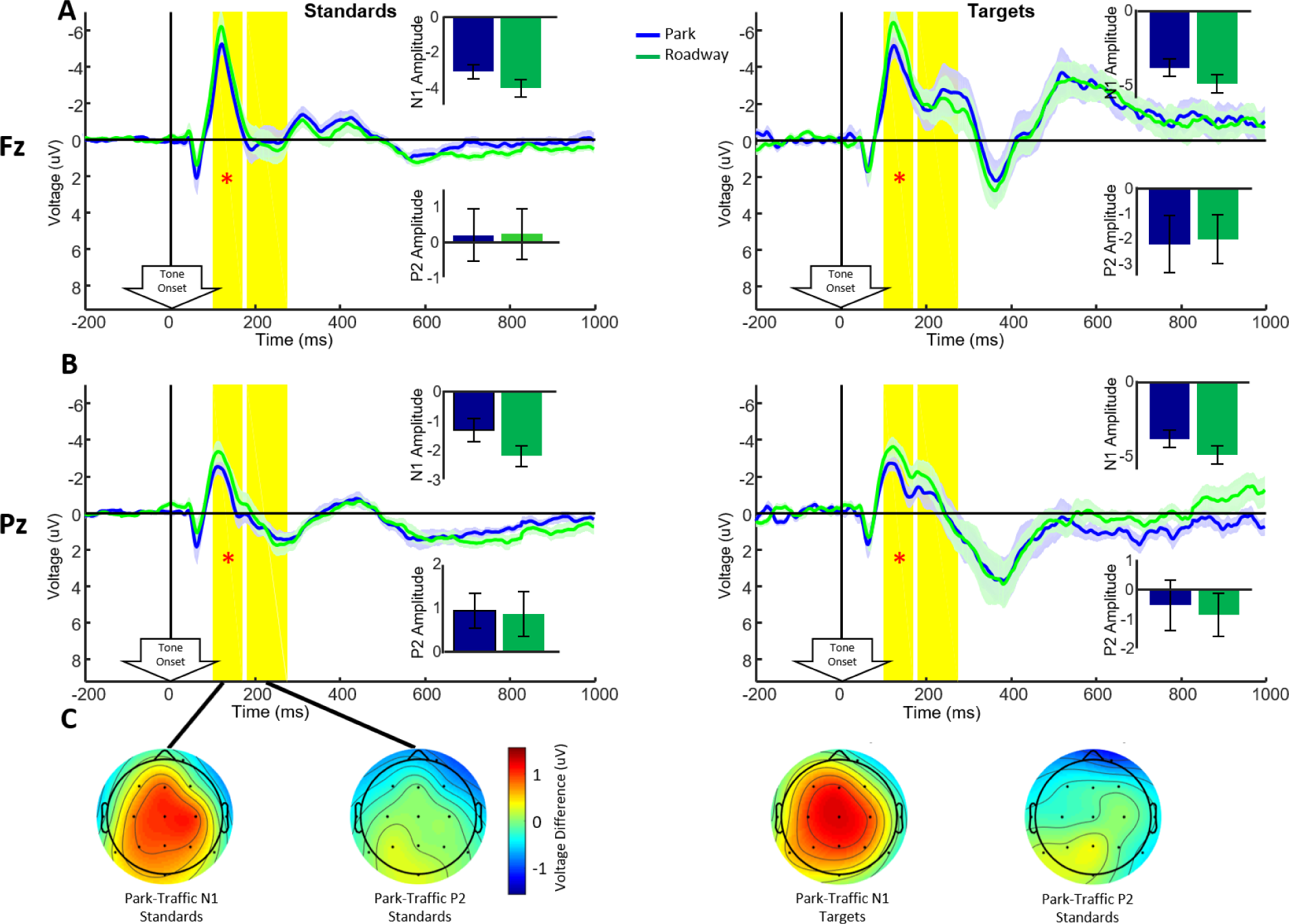
Grand average ERPs A: Grand average ERPs collected at the Fz electrode location, plotted separately to compare standards and targets between conditions. Shaded regions indicate the within-subjects standard error of the mean. Inset bar graphs show the mean and within-subjects standard error across participants in the N1 and P2 time windows. B: Grand average ERPs for the Pz electrode location, plotted separately to compare standards and targets between conditions. Inset bar graphs depict mean and within-subjects standard error across participants in the N1 and P2 time windows. C. Topographies of the N1 and P2 time window plotted for the park-roadway condition differences.

**Table 1.**
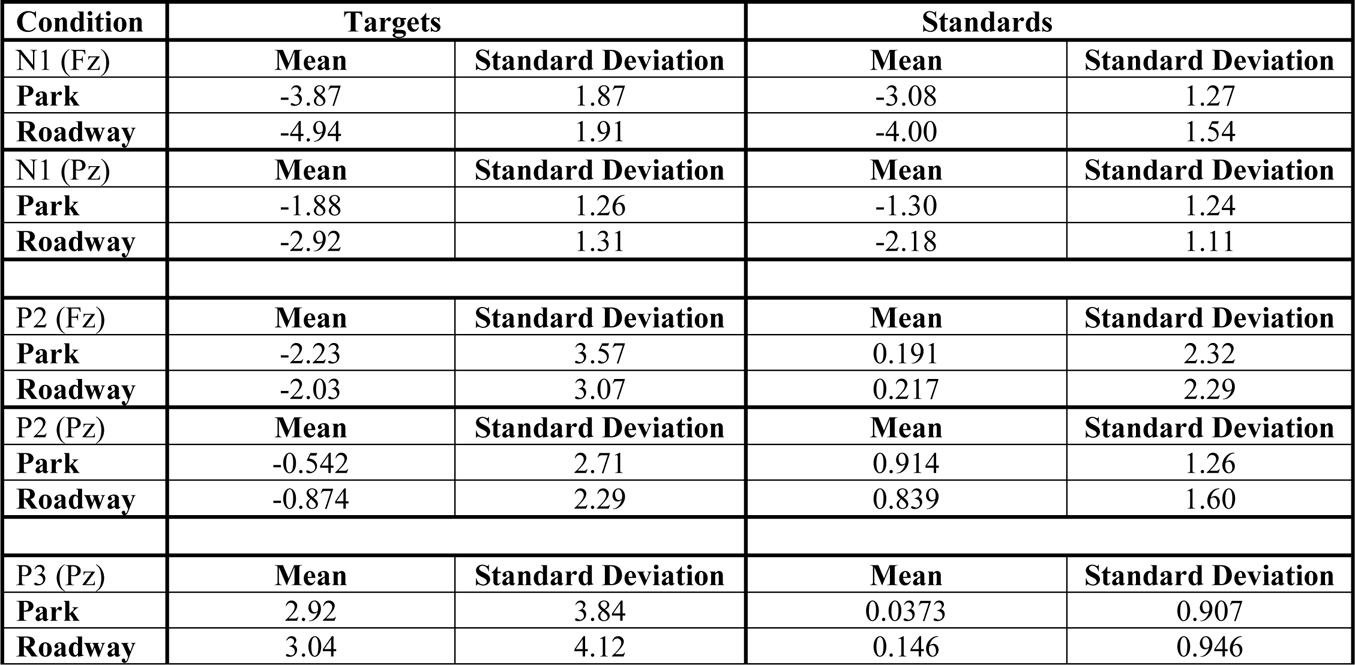
Mean and standard deviation of N1, P2 and P3 ERP components for both standard and target stimuli.

| Condition | Targets |  | Standards |  |
| --- | --- | --- | --- | --- |
| N1 (Fz) | Mean | Standard Deviation | Mean | Standard Deviation |
| Park | -3.87 | 1.87 | -3.08 | 1.27 |
| Roadway | -4.94 | 1.91 | -4.00 | 1.54 |
| N1 (Pz) | Mean | Standard Deviation | Mean | Standard Deviation |
| Park | -1.88 | 1.26 | -1.30 | 1.24 |
| Roadway | -2.92 | 1.31 | -2.18 | 1.11 |
| P2 (Fz) | Mean | Standard Deviation | Mean | Standard Deviation |
| Park | -2.23 | 3.57 | 0.191 | 2.32 |
| Roadway | -2.03 | 3.07 | 0.217 | 2.29 |
| P2 (Pz) | Mean | Standard Deviation | Mean | Standard Deviation |
| Park | -0.542 | 2.71 | 0.914 | 1.26 |
| Roadway | -0.874 | 2.29 | 0.839 | 1.60 |
| P3 (Pz) | Mean | Standard Deviation | Mean | Standard Deviation |
| Park | 2.92 | 3.84 | 0.0373 | 0.907 |
| Roadway | 3.04 | 4.12 | 0.146 | 0.946 |

Additionally, as previous studies have shown increased background noise to decrease the amplitude of the P2 component between 175 and 275 ms, one may expect to observe a similar effect in which the roadway condition shows a lower P2 amplitude. However, evident from the plots in Figure 6A and B, no difference in P2 amplitude exists between the two conditions.

Again, 2 × 2 repeated measures ANOVA tests were carried out at the Fz and Pz electrode locations, showing a significant effect of stimulus (Fz: *F*(1,9) = 13.2, *p* = 0.0054, η_p_^2^ = 0.595; Pz: *F*(1,9) = 5.71, *p* = 0.0405, η_p_^2^ = 0.388), with higher amplitude for standards than targets (Table 1). No effect was found for the condition (Fz: *F*(1,9) = 0.0797, *p* = 0.784, η_p_^2^ = 0.0090; Pz: *F*(1,9) = 0.256, *p* =0.625, η_p_^2^ = 0.028) or interaction (Fz: *F*(1,9) = 0.154, *p* = 0.704, η_p_^2^ = 0.017; Pz: *F*(1,9) = 0.422, *p* = 0.532, η_p_^2^ = 0.045).

### 3.7 Combined experimental analyses

Figure 7A and B shows two bar graphs of N1 and P2 amplitude, respectively, compared across 4 experiments which all used the oddball task (Scanlon et al., 2017a; Scanlon et al., 2017b; Scanlon et al., 2019; Current study). All of these experiments used the same auditory oddball task. All of these experiments were performed on different participants. Experiment 1 is a previous study in which participants performed an auditory oddball task on a stationary bicycle (Scanlon et al., 2017a). While performing the oddball task, participants were asked to sit still (pre), pedal sub-aerobically (bike) and sit still again (post). Experiment 2 is a previous study in which participants performed the oddball task both while sitting inside a laboratory faraday cage (inside) and cycling outside the lab (outside; Scanlon et al., 2017b). In experiment 3, participants performed the auditory oddball task with four different background noise and volume conditions (Scanlon et al., 2019). The task was performed with the presence of background outdoor sounds recorded near a roadway (sound), with the presence of background white noise (white), with a silent background and the tones played at reduced volume (silent low), and with a silent background and regular tone volume (silent). Experiment 4 is the current study, with the oddball task performed in the quiet park (park) and next to the noisy roadway (roadway). Figure 7A shows the N1 amplitudes to standard tones in 11 conditions from four experiments using the same auditory oddball task. Evident from the graph is that experimental conditions in which the participant performed the task in non-ideal auditory conditions (e.g. with background noise or low volume; shades of green and yellow) show generally larger N1 amplitudes than when the task was performed in ideal situations (silent background, normal volume; shades of grey).

**Figure 7.**
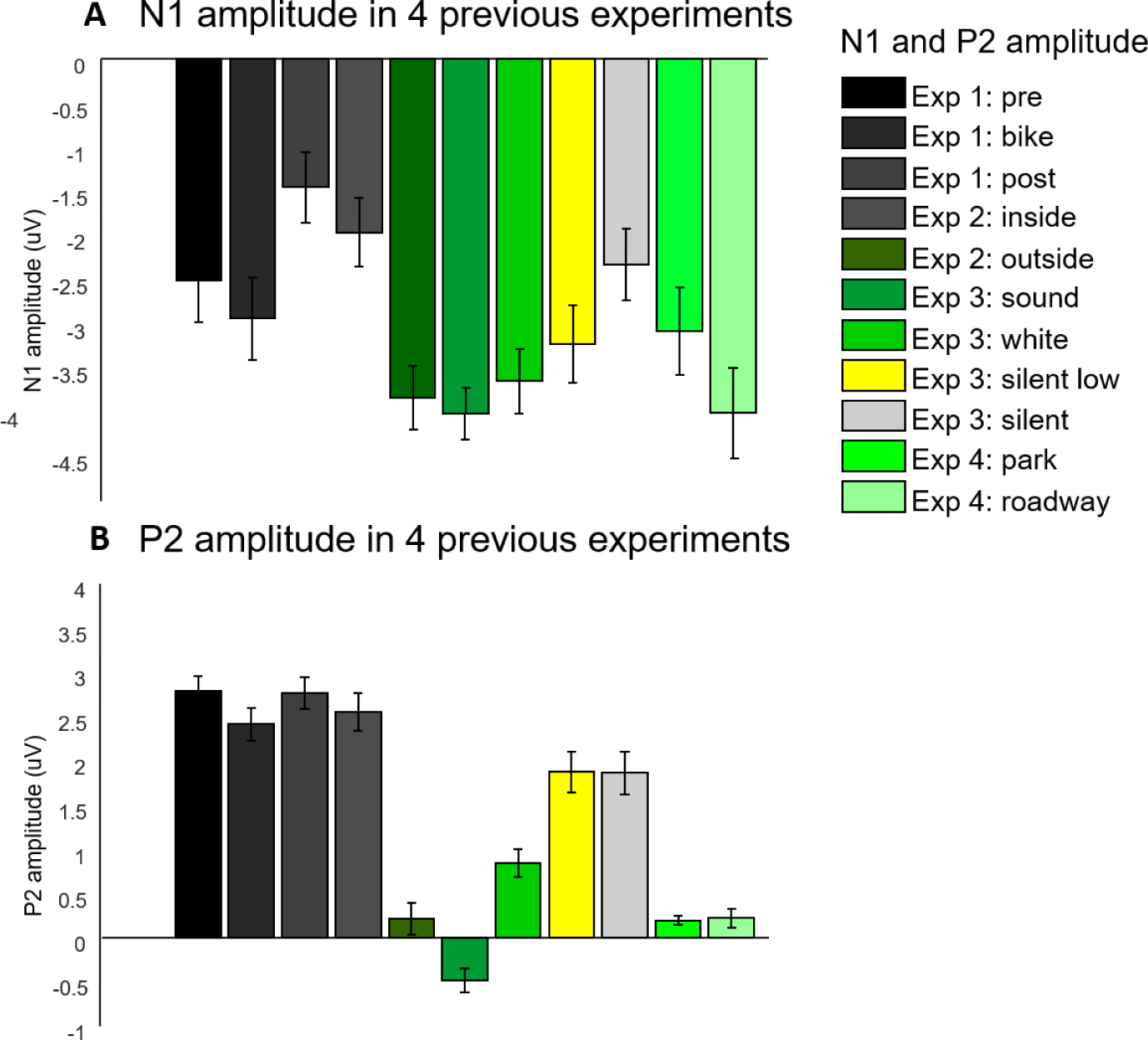
Combined experimental comparisons of the N1 and P2 amplitude. A: Bar graph of mean N1 amplitude to standard tones in 11 conditions across four auditory experiments using the same auditory oddball task. Shades of green are used to represent experimental conditions with background sounds, and shades of grey are used to represent conditions with silent backgrounds. Yellow is used to depict the low-volume tones condition of experiment 2. Error bars depict the within-subjects standard error of the mean. B: Bar graph of mean P2 amplitude to the standard tone in 11 conditions across four auditory experiments using the same auditory oddball task. Shades of green are used to represent experimental conditions with background sounds, and shades of grey are used to represent conditions with silent backgrounds. Yellow is used to depict the low-volume tones condition of experiment 2. Error bars depict the within-subjects standard error of the mean. Experiment references: Exp 1: (Scanlon et al., 2017a), Exp 2 (Scanlon et al., 2017b), Exp 3 (Scanlon et al., 2019), Exp 4 (Current study).

Figure 7B depicts the P2 amplitudes to standard tones in 11 conditions from four experiments using the same auditory oddball task. The graph shows that experimental conditions in which the participant performed the task with background noise (shades of green) have generally lower P2 amplitude than those in which the task was performed in silence (shades of grey and yellow).

## 4. Discussion

In this study we examined the effect of two different environments on brain activity, by having participants perform the same auditory task both next to a noisy roadway and in a quiet park. We chose the task to be an auditory oddball task while riding a bicycle at a slow sub-aerobic pace, in order for it to be both similar to a real-world task, and for the data to be comparable to previous studies within our lab. Our main findings showed that performing the task in a noisier environment changed the way participants processed the task, as demonstrated by an increase N1 amplitude.

### 4.1 Behavioural differences

As in previous studies (Scanlon et al., 2017a; Scanlon et al., 2017b; Scanlon et al., 2019; Scanlon et al., *In preparation*), the ERP differences between conditions did not appear to contribute to behavioural differences in the oddball task, as there were no significant differences in reaction time between conditions. This indicates that the N1 may reflect a compensatory process, allowing the mind to put greater focus on a target when distractions are present. There was however a nearly significant difference in reaction accuracy between the two conditions (p=0.0729) with a moderately large effect size (*Cohen’s d* = −0.642). This indicates a possibility that this difference could be significant with a larger sample size, and possibly point to a behavioural link with modulations in the N1. This should be explored further in future research.

### 4.2 Data noise

As this study took place outdoors and we had little control over the environment itself, we considered it important to measure the effect the environment might have on our EEG data noise. Figure 3 demonstrates that the park condition had a larger amount of baseline noise than the roadway condition (Figure 3), with higher RMS in the baseline EEG period for both trial averaged ERP and single-trial data. This may have possibly been due to slightly higher levels of elevation change within the park. While we were careful to find two environments that were similar in elevation, real world studies often have the constraint of being unable to change what the current study can access in the real world. However, this increase in data noise was relatively small compared to differences between stationary and mobile recording situations in previous studies (Scanlon et al., 2017b), in which the ERPs recorded appeared largely unaffected by this difference.

### 4.3 Spectral Power

Figure 4A demonstrated increased beta power (15-30Hz) at Fz in the park condition, possibly due to the increase in mechanical noise, similar to increased high-frequency oscillations in the beta range found in previous studies during stationary pedaling (Scanlon et al., 2017a) and outdoor biking (Scanlon et al., 2017b) compared to sitting still. Alternatively, this increase in beta power could be related to changes in pedalling effort, due to increased elevation changes in the park condition, as movement initiations such as bicycle pedalling have been shown to alter beta power in previous studies (Storzer et al., 2016; Jain et al., 2013). However, the current design is unable to disentangle these possibilities.

### 4.4 P3 morphology

As expected there were no significant differences observed between the two conditions for the P3 component. As the participants were performing the same dual task in both conditions (i.e. cycling during the oddball task), and it is known that dual tasks tend to reduce P3 amplitude, it appears that the P3 component was equally reduced between the two conditions. This is evident because the peak amplitude for the P3 for both conditions was approximately 2 μV lower than those observed in previous studies done by this lab in which there was no concurrent task (Scanlon et al., 2017a; Scanlon et al., 2017b). Given the well-established observation that performing a secondary task during an auditory oddball task will reduce cognitive resources available for the task, therefore decreasing target-standard P3 amplitude (Wicken et al., 1983; Polich, 1987; Kramer & Strayer, 1988; Polich & Kok, 1995; Scanlon et al., 2017b), this observation was not unexpected.

### 4.5 N1 and P2 morphology

Previous studies indicated that ambient noise during an oddball task decreased the P2 and increased N1 amplitude for both standards and targets during the auditory oddball task due to a process of filtering out irrelevant sounds in order to perform the task (Scanlon et al., 2017b; Scanlon et al., 2019). We initially expected that we would find a similar difference in this study, with a reduced P2 and increased N1 while participants were cycling next to the noisy roadway compared to the quiet park. However this was not the case, as no significant differences were found for the P2 component. This may indicate that the excess noise of the roadway did not sufficiently create the necessity to filter out ambient noise, or, that this noise filtering process was present during both conditions. If one compares the current study to Scanlon et al. (2017a) or Scanlon et al. (2017b), the P2 amplitude in both current conditions appears to be at least 2 μV smaller than the indoor conditions of the previous studies. This leads us to deduce that the lower amount of ambient noise in the park condition was sufficient for the P2 to be reduced to an equal degree as the roadway condition, and therefore this effect was observed equally in both conditions. In contrast to this however, there was a significant difference in the N1 between the two conditions with a larger N1 amplitude while participants performed the task near the roadway. This is consistent with the increase in ambient noise shown in previous studies. This deviation indicates that while the N1 and P2 had previously been assumed to be modulated simultaneously to represent the same process, this may not be the case. Here, it appears that the level of ambient background noise had the ability to augment the N1, but any amount of ambient noise is sufficient to reduce the P2.

The N1-P2 complex is believed to be reflective of pre-attentive sound processing within the auditory cortex (Näätänen & Picton, 1987). The components of this complex, including the P1, N1 and P2 have been demonstrated to relate to several temporally overlapping processes which originate near or within the primary auditory cortex (Näätänen & Picton, 1987; Wolpaw and Penry, 1975; Wood & Wolpaw, 1982). P2 in particular has been shown to relate to cognitive functions such as working memory, memory performance and semantic processing during contextually based tasks (Dunn et al., 1998; Federmeier & Kutas, 2002; Lefebvre et al., 2005).

The P2 is also thought to reflect a component of top-down cognition and perceptual processing, and may represent a process of inhibiting one’s perception of unimportant or repetitive stimuli in order to perform a task (Luck & Hillyard, 1994; Freunberger et al., 2007). Additionally, the P2 has been hypothesized to reflect a process of suppressing the perception of irrelevant stimuli to allow stimulus discrimination within a primary task (Potts et al., 1996; Potts, 2004; Kim et al., 2008; Getzmann et al., 2016). The auditory P2 appears to relate to the subjective difficulty of stimulus discrimination, as both the auditory P2 and N1-P2 complex have been reliably found to have increased amplitude following discrimination training (Atienza et al., 2002; Hayes et al., 2003; Reinke et al., 2003; Trainor et al., 2003; Tremblay et al., 1997, 2002, 2001). More specifically to our task, one study showed a decrease in auditory P2 amplitude during a speech discrimination task while participants had to ignore irrelevant background speech in a ‘cocktail party’ scenario, compared to a condition with no background distractions (Getzmann et al., 2016). This could explain why the P2 appears to have been reduced in both conditions, as the auditory P2 appears to be reduced any time stimulus discrimination in the task is more difficult, and therefore any amount of ambient noise could have this effect.

While the N1 and P2 are clearly related in a cluster of processes that take place within the N1-P2 complex, they appear here to serve similar but distinct functions. Previous research has proposed that the N1 itself appears to reflect several different auditory processes, with up to six distinct components (Näätänen & Picton, 1987). The most alterable of these being influenced by several contextual factors including a ‘sensory acceptance-rejection’ factor, which appears to attenuate all responses to uninteresting, irrelevant or unpleasant sensory inputs while enhancing responses to pleasant, interesting or important stimuli. The N1 tends to appear with shorter latency and larger amplitude when the attended and unattended sounds can be easily distinguished by physical cues such as pitch or location (Näätänen, 1982, 1992). For example, the N1 has been shown to increase when an auditory stimulus occurs in the location where an individual is attending, (Teder-Sälejärvi, et al. 1998) to allow an individual to enhance the auditory perception of this stimuli. This effect is increased in individuals who are blind (Roder et al., 1999). Overall, N1 amplitude is said to increase as a function of increased attentional allocation to a particular auditory input channel, which correlates to increased behavioural accuracy for targets in that channel (Hink, Voorhis, Hillyard, & Smith, 1977). Functionally, the amplitude and latency of the N1 is believed to represent the quantity of sensory information moving through an early auditory channel selection mechanism (Hillyard. Hink, Schwent, & Picton, 1973), greater processing of attended channel information (Okita, 1981; Näätänen, 1982), as well as how well-matched the eliciting stimulus and cue characteristics are within the attended auditory input channel (Näätänen, 1992). In this study, it appears that while participants had to filter out noise in both conditions, the roadway condition required a larger amount of attention and processing of the task tones in order to perform the task, and therefore the N1 was increased.

### 4.6 Limitations

Similar to our previous studies (Scanlon et al., 2017a; Scanlon et al., 2017b), slow cycling was used here in order to create a mobile task with minimal head movement or aerobic effects. This experimental setup also has some drawbacks in that it required several items to be carried around with the participant in the backpack. Further, the amplifier used was bulky and required long electrode wires to reach from the head to the backpack. Recent advances with EEG systems which are wireless and made specifically for mobile EEG (Debener et al., 2015; Zink et al., 2016; Hashemi et al., 2016; Krigolson et al., 2017) would greatly improve the portability of our system. We have recently started testing the use of miniature computer systems, such as the Latte Panda, however testing these new technologies also lead to some technical problems in this experiment. These technical problems included decreasing strength in the Anker astro battery, which lead to power loss in our original recording computer (Latte Panda). This required us to use instead the Microsoft surface, and discard subjects whose data was successfully recorded on the first computer. Additionally a few datasets had malfunctions in connections between the Raspberry Pi and the recording computer, leading to loss of data triggers. In the future we plan to continue testing new technologies while streamlining our current devices in order to avoid these issues in the next studies.

An additional limitation exists within the sample size of 10 subjects, which is relatively low. This low sample size is due to some technical problems (see above paragraph) that arose during data collection, as well as the limited amount of time in the season available to record outside in Edmonton. However, given the high level of within subject precision due to high number of trials with consistent patterns, classic measures of power are less appropriate to estimate reliability of results (Luck, 2019). The study is based on similar previous studies with a sample size of 12 (Scanlon et al., 2017b) and 14 (Scanlon et al., 2017a), which both had comparable amounts of data noise due to movement conditions and were able to show results within the ERP components selected for this study. Therefore, we believe that this study has a similar ability to present accurate results.

A final limitation exists in that we did not counterbalance the tones in our oddball task. While non-counterbalanced oddball tasks are common in the literature (Ladouce, et al., 2019; Mathewson et al., 2015; Laszlo, et al., 2014; Zink, et al., 2016), this does leave the possibility of creating problems for participants with hearing problems that they were not previously aware of (as we did not take participants with known hearing problems), as higher pitched sounds are often lost first during aging (Gates & Mills, 2005). There is also a possibility for an unequal interaction between the different tones and the background sounds, however because we found the same effect for both standards and targets between the conditions, it does not appear that this was an issue. Nonetheless we see the advantage to counterbalancing oddball tones and plan to take up this practice in future studies.

### 4.7 Combined experimental comparisons

Figure 7A and B depicts the main measures (i.e. the N1 and P2 amplitude) compared across 4 experiments from our lab which all used the oddball task (Scanlon et al., 2017a; Scanlon et al., 2017b; Scanlon et al., 2019; Current study). Figure 7A demonstrates that experimental conditions in which the participant performed the task in non-ideal conditions (i.e., with background noise or low volume) had a higher N1 amplitude than conditions in which the task was performed in ideal conditions (i.e. silent background with normal volume). This reinforces the effect of attention on N1 amplitude, as non-ideal conditions require the participant to ‘tune in’ to the stimulus more than they might have to in ideal conditions. Figure 7B shows that experimental conditions in which the participant performed the task in the presence of background sounds had generally lower P2 amplitude than conditions in which the task was performed in silence. This helps to demonstrate the way the P2 represents ‘tuning out’ distracting sounds in auditory tasks. The dichotomy between the functions of the N1 and P2 can especially be seen in the silent-low condition of experiment 2, as this condition had an enhanced N1 with a non-affected P2, demonstrating that the low-volume task required enhanced attention, with no requirement to tune out irrelevant information. The two conditions of the current study also demonstrate this dichotomy, as both of these conditions have a larger N1 amplitude than any of the conditions performed in an ideal auditory situation, and the roadway condition has a significantly higher N1 than the park condition. However the P2 amplitude for both of these outdoor conditions is near zero and much lower than any condition in which the task was performed in silence, demonstrating that while there was no P2 difference within this experiment, the P2 was likely reduced in both conditions due to background sounds in the outdoor environment.

### 4.8 Future directions

This research program had the ultimate goal of measuring ERPs which naturally occur in the real world, in order to gain a deeper understanding of how the brain functions during everyday life. We have recently begun testing EEG set-up which uses video capture and coding to identify moments in the EEG data when interesting events happen in an individual’s environment. We have also started to test the use of virtual reality (VR) to create environments and allow stimulus presentation, which also opens up the possibility of testing the brain in any unlimited number of simulated environments.

### 4.9 Conclusions

In this study we used a fully mobile EEG task to show that the environment in which one performs a task is enough to alter brain activity. We also demonstrated that the N1 and P2, while functionally related, clearly represent two separate processes in the early auditory system. It appears that the N1 increases when a situation requires increased attentiveness to a stimulus, while the P2 is altered any time the auditory task is more difficult to discriminate. As shown in previous studies, this effect appears to be independent of movement and behavioural measures.

**6. Supplementary Materials**

